# A fitness distribution law for amino-acid replacements

**DOI:** 10.1101/2024.10.11.617952

**Authors:** Mengyi Sun, Arlin Stoltzfus, David M. McCandlish

## Abstract

The effect of replacing the amino acid at a given site in a protein is difficult to predict. Yet, evolutionary comparisons have revealed highly regular patterns of interchangeability between pairs of amino acids, and such patterns have proved enormously useful in a range of applications in bioinformatics, evolutionary inference, and protein design. Here we reconcile these apparently contradictory observations using fitness data from over 350,000 experimental amino acid replacements. Almost one-quarter of the 20 *×* 19 = 380 types of replacements have broad distributions of fitness effects (DFEs) that closely resemble the background DFE for random changes, indicating an overwhelming influence of protein context in determining mutational effects. However, we also observe that the 380 pair-specific DFEs closely follow a maximum entropy distribution, specifically a truncated exponential distribution. The shape of this distribution is determined entirely by its mean, which is equivalent to the chance that a replacement of the given type is fitter than a random replacement. In this type of distribution, modest deviations in the mean correspond to much larger changes in the probability of falling in the far right tail, so that modest differences in mean exchangeability may result in much larger differences in the chance of a highly fit mutation. Indeed, we show that under the assumption that purifying selection filters out the vast majority of mutations, the maximum entropy distributions of fitness effects inferred from deep mutational scanning experiments predict the characteristic patterns of amino acid change observed in molecular evolution. These maximum entropy distributions of mutational effects not only provide a tuneable model for molecular evolution, but also have implications for mutational effect prediction and protein engineering.

## Introduction

Since the earliest studies of homologous amino acid sequences in the 1960s, it has been clear that amino acid replacements often follow highly regular patterns (1, 2). For instance, threonine and serine replace each other much more frequently than most pairs of amino acids. One straightforward interpretation for these patterns is that some pairs of amino acids are more similar to each other in terms of physicochemical properties, and thus are functionally more interchangeable (1). This way of thinking brought attention to diverse ways of measuring the physicochemical similarity of amino acids (3), and led to the development of composite measures such as Grantham and Miyata distances (4–7). Over time, cost matrices reflecting the evolutionary replaceability of amino acids became an indispensable part of how we align proteins and search sequence space (8–10). What these approaches imply is that a particular type of replacement has a characteristic effect determined by the similarity of the amino acids.

Yet many other lines of evidence suggest that the effect of an amino acid replacement depends sensitively on details of the structural context (11–13), to the point that the identities of the wild-type and mutant amino acids, taken in isolation, are relatively uninformative. Broad evidence supporting this interpretation includes the difficulty in reliably predicting the pathogenicity of individual SNPs (11), and the detailed biophysical analysis typically required to understand the effects of individual amino acid replacements on protein function (14). Context-dependence has been widely demonstrated in the literature on genetic epistasis (15), the phenomenon whereby the state at one genetic site influences the effects of changing a second site. Such epistatic interactions are common among beneficial mutations during natural adaptive evolution (16) and during protein engineering (17), as well as among the substitutions accumulated during long-term molecular evolution (18, 19).

In an attempt to reconcile these two apparently contradictory lines of evidence, here we aggregate data across many different proteins (20, 21) to construct a distribution of fitness effects (DFE) for each of the 380 possible types of amino acid replacement using data from deep mutational scanning assays (22, 23). Such an approach has become possible due to advances in high-throughput mutagenesis and quantification by deep sequencing, allowing thousands of individual amino acid mutations to be assayed in a single study; we can then aggregate data across studies by changing the measured values to percentile ranks.

We find that, for a large fraction of replacement types, the type-specific distribution of fitness effects (DFE) is not substantially different from the overall background DFE. Even the types of replacements that are most likely to be benign, or most likely to be deleterious, have wide distributions of fitness effects, consistent with a strong role of molecular context in determining mutational effects.

However, we also observe that once converted from the original measurement scale into percentile ranks, these type-specific distributions of mutational effects share a common functional form. Specifically, the type-specific distributions take the form of the probability distribution with maximum entropy given the mean effect of a particular type of substitution, which in our context results in a truncated exponential distribution. These truncated exponential distributions have the feature that differences in the probability density of effects are most extreme at the tails. Because of this, two distributions that differ only modestly in the mean value may show a large difference in the chance of being among the most benign mutations, i.e., at the top end of the fitness distribution. This sensitivity in the tails suggests an explanation for why evolutionary divergence shows strong preferences for specific amino acid replacements despite the broad distributions of effects observed for most types of exchanges. We test this idea using a model of protein sequence change under purifying selection, where only the most benign mutations are allowed to fix. We show that this type of model largely accounts for patterns of amino acid replacement in molecular evolution. This finding suggests a resolution for the apparent conflict in the predictability of amino-acid replacements, with implications for understanding molecular evolution, advancing protein engineering, and addressing diseases triggered by amino acid changes. The universal form we observe for type-specific DFEs also suggests a way to interpret distributions for other types of mutant effects, i.e., other than fitness.

## Results

### Fitness data from deep mutational scanning studies

To uncover the patterns of DFEs for 380 ordered pairs of amino acids, we leveraged the ProteinGym dataset (24), which encompasses about 350,000 single missense variants derived from 87 distinct deep mutational scanning (DMS) assays. These assays cover a wide range of functional properties, including cellular growth rates, ligand binding, and drug resistance, and encompass 58 distinct protein families from diverse taxa (45 eukaryotes, 21 prokaryotes, and 21 viruses). ProteinGym aims to establish measurement benchmarks for mutation effect prediction in machine learning algorithms that utilize evolutionary information. Due to its large size and its emphasis on DMS assays that directly measure growth phenotypes (48 out of 87 assays) or protein properties closely related to fitness, ProteinGym is well suited to investigate DFEs systematically.

To combine the results of different experiments, which may have different measures and scales of effect, we rank the mutant effect scores from each experiment and transform them into quantiles, which vary between 0 and 1. When scores are transformed in this way, the distribution of effects over all changes is uniform, and when a specific type of change (e.g., Ala to Val) has an average score of *x*, this means that *x* is the chance that a mutation of this type is more fit than a randomly chosen mutation. Using this approach, we constructed 380 distinct DFEs, each defined within the range of [0,1]. The number of data points for each DFE varies, with an average of about 942.3 replacements per type (median 1004), and a range from 265 for the Trp→Ser replacement to 1818 for Leu→Pro.

### DFEs for 380 types of amino acid changes

What shapes do we expect for the DFEs for replacing one amino acid with another? If the replacement effect is mostly determined by the physicochemical similarity between two amino acids, as suggested by patterns of sequence evolution, we would expect the DFE for a particular amino acid pair to be very sharp. For instance, for amino acid pairs that are highly dissimilar, the probability mass would be concentrated near 0 (Fig. 1, top left); for moderately dissimilar amino-acid pairs, the probability mass would have an intermediate value (Fig. 1, top middle); and for very similar amino-acid pairs, the probability mass would be concentrated near 1 (Fig. 1, top right). By contrast, if context effects dominate, we might see a much flatter DFE, e.g., the extreme case of a uniform distribution over the interval [0,1] (Fig. 1, bottom left) indicates that the DFE for replacements of a given type is identical to the overall DFE from the same experiment. One also could imagine more complex situations, such as replacements that are benign in one background but deleterious in another (e.g., surface vs. buried), resulting in a bimodal distribution (Fig. 1, bottom right).

**Fig. 1.**
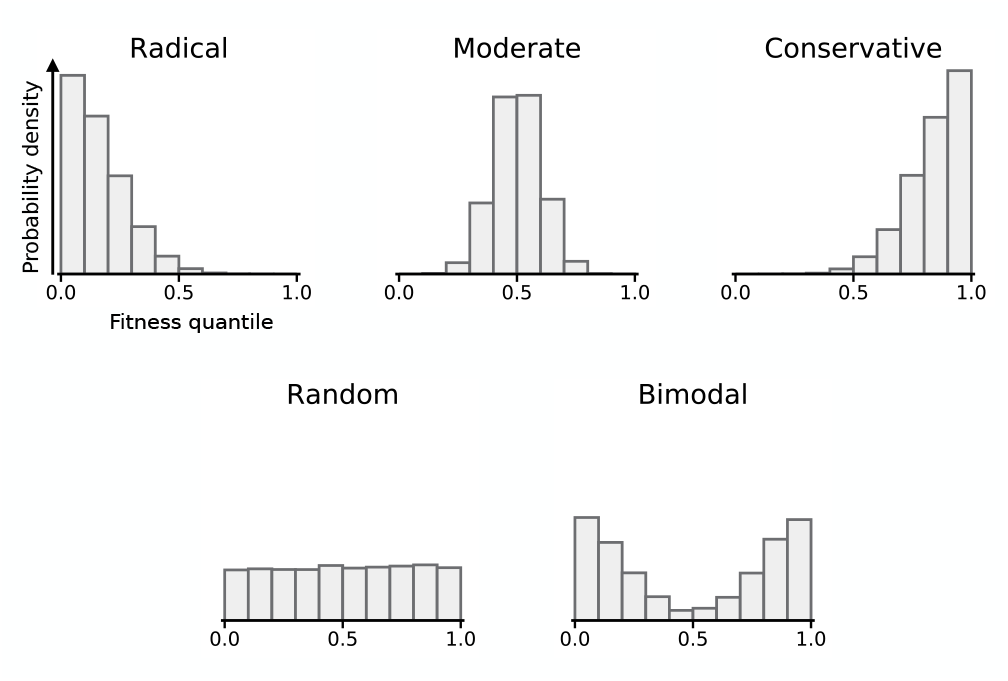
Illustration of type-specific DFEs as quantile distributions. Each histogram shows the probability that a replacement of a given type will lie at a given fitness quantile relative to the effects of other amino acid replacements in the same protein. A distribution that is concentrated at low values indicates that the corresponding type of replacement tends to be deleterious, whereas types of substitution that tend to be benign will have distributions concentrated at high values. Mutational types that have the same DFE as random mutations will appear as uniform distributions.

How do the actual distributions compare to these expectations? Fig. 2 shows the DFEs for all 380 types of amino acid changes. None of the distributions are particularly narrow; instead, each distribution spreads broadly over the whole set of 15 quantile bins (note that this is also true for the marginal distributions aggregated by the “from” or “to” amino acid, shown in Fig. S1). If we use the Kolmogorov–Smirnov statistic to compare these distributions with uniform distributions, where the amino-acid identities are maximally uninformative, we find that most have only a small difference from the uniform distribution (Fig. S2a, median KS-statistic=0.13, where the KS-statistic is given by the maximal absolute difference between the empirical cumulative distribution function and the cumulative distribution function under the null). Close to one-quarter of the distributions (92 out of 380) are statistically indistinguishable from a uniform distribution (Bonferroni-corrected *p >* 0.05, Kolmogorov–Smirnov test).

**Fig. 2.**
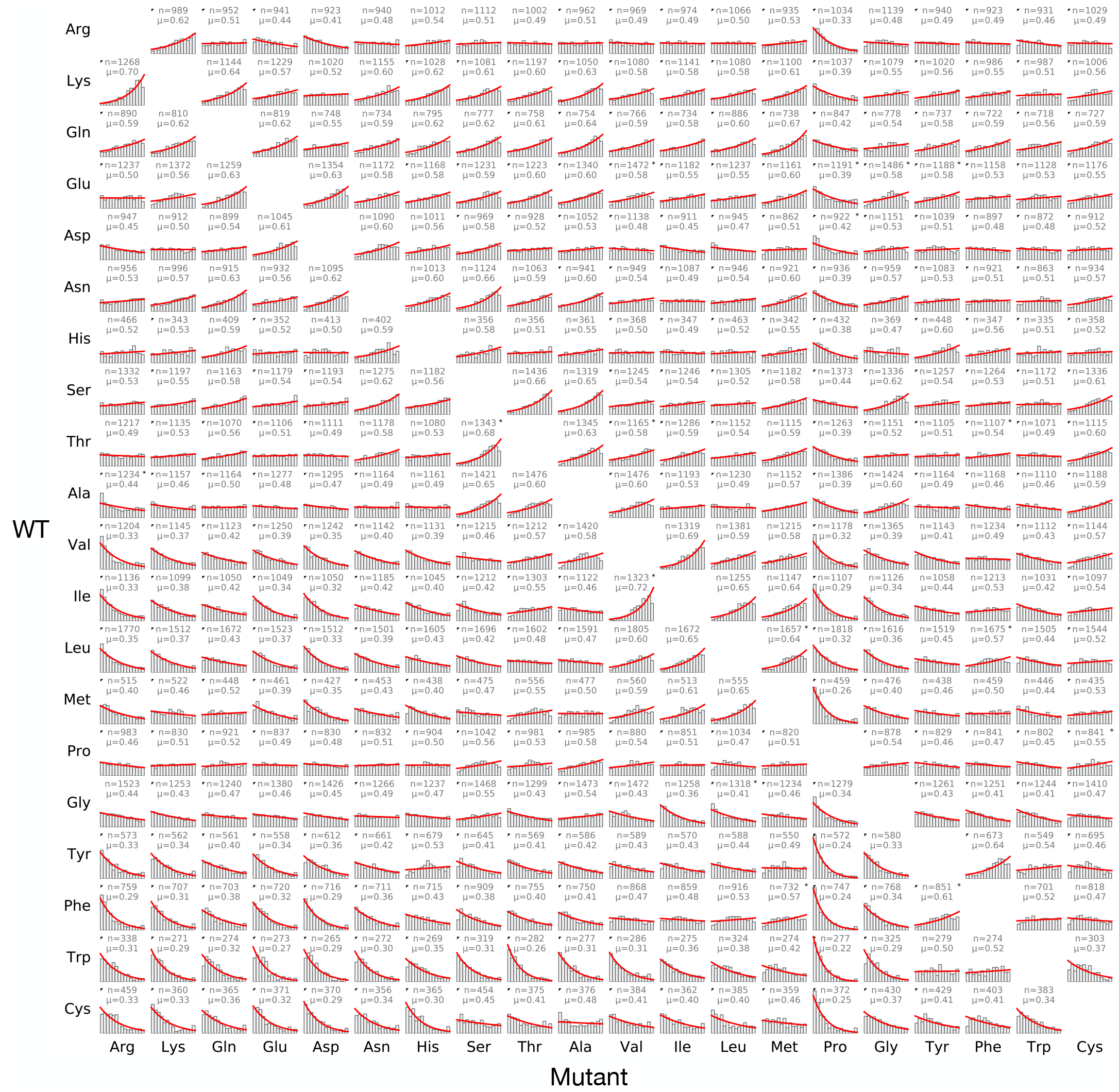
Distributions of fitness effects for 358,080 amino-acid-altering mutations, categorized by the wild-type (row) and mutant (column) amino acids. Histograms show fitness quantiles observed for each type of amino-acid replacement, along with the corresponding truncated exponential fit (red lines). For each distribution, *μ* is the mean (equal to the probability that a replacement of this type is fitter than a random mutation) and *n* is the number of observations. Distributions that deviate significantly from a truncated exponential distribution are marked with a star at the right corner, whereas black triangles in the upper left corner indicate the pairs for which forward-reverse asymmetry is significant (Bonferroni-corrected *p <* 0.05, two-sample Kolmogorov–Smirnov test).

For the distributions that are statistically distinct from a uniform distribution, another interesting pattern emerges: none of these distributions appear to be strongly bimodal, rather the distributions tend to be highly monotonic, i.e., the density of mutations in a bin either increases consistently from low to high, or decreases consistently. This monotonicity can be quantified by the absolute value of the Spearman correlation between the rank order of each bin and its size (i.e., counts), with the result that the correlations tend to be high, with a median of 0.91 (Fig. S2b). Finally, one subtle prediction from the physicochemical distance model illustrated in the top row of Fig. 1 relates to the fact that distances are symmetrical, so that the biochemical distance between amino-acid *i* and amino-acid *j* is the same as the distance between amino-acid *j* and amino-acid *i*. Thus, in the absence of context effects, the distributions for *i → j* and *j → i* should be the same. However, this symmetry does not hold for 134 out of 190 pairs of amino acids (distributions marked with triangles; Bonferroni-corrected *p >* 0.05, two-sample Kolmogorov–Smirnov test). This kind of asymmetry again indicates the importance of molecular context.

### A universal statistical law governing the 380 DFEs

Motivated by the qualitative similarities between the shapes of the 380 DFEs, we sought to determine whether all of these DFEs could be explained as belonging to a single family of probability distributions. Maximum entropy probability distributions arise in many areas of statistics and the natural sciences and are defined as distributions that maximize the entropy (intuitively, the uniformity of the distribution), subject to certain constraints (25–27). For example, the normal distribution is the maximum entropy distribution over the real numbers with a specified mean and variance and the geometric distribution is the maximum entropy distribution over the natural numbers with a specified mean. Maximum entropy distributions have also been applied successfully in various biological contexts, including modeling the distribution of mutational effects on Malthusian fitness (28), site-specific amino acid distributions (29), and cell packing geometries (30). For our purposes, the constraints on a type-specific DFE for quantiles are (1) that it is a distribution on the interval [0,1] and (2) that the distribution has a particular mean, which in this case is also equal to the probability that a mutation of that type is fitter than a random mutant. In principle, we could consider higher-order moments like variance and skewness as additional constraints, but importantly, when the mean is plotted against the variance for the 380 DFEs, the points align along a one-dimensional curve (Fig. S3a, red points), indicating that the variance itself appears to be a function of the mean. Consequently, we constrain our distributions up to only their first moments (mean).

Under these constraints on the domain of the probability distribution and its mean, the maximum entropy distribution turns out to be a truncated exponential distribution (27) which can be expressed as:

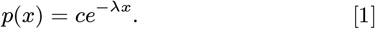

Here, 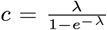 is a normalizing constant, and *λ* is a real-valued parameter. This distribution has a density that is either monotonically increasing or decreasing or else constant (when the mean is 0.5), consistent with our observations of the 380 DFE shapes. Furthermore, this family of distributions predicts a very specific mean-variance relationship, which is well-matched by our data (Fig. S3a).

To further evaluate how well these distributions fit the 380 DFEs, we estimated *λ* using the means of each DFE (see Materials and Methods). To avoid overfitting, for each DFE we used half the data to estimate the mean, and the other half to test the deviation from the estimated truncated exponential distribution. As can be shown in Fig. 2, in over 95 % of cases, the observed DFEs were statistically similar to the estimated ones (Bonferroni-corrected *p >* 0.05, Kolmogorov–Smirnov test), with generally small KS-statistics (Fig. S3b).

Different DMS experiments may have systematically different distributions of the 380 possible replacements (due to differing amino acid composition of the target protein and different library construction methods). To probe whether such differences might systematically influence shapes of DFEs, we focused on 28 datasets that cover all types of replacements, equalizing representation of each mutation type by either up-sampling to the maximum or down-sampling to the minimum count. We found that the *λ* values from these adjusted datasets closely matched those from all 87 datasets (Fig. S3c and d).

Whereas the exponential distribution law fits well to fitness quantiles, it does not necessarily apply to the quantile distributions for other measures of mutational effect from the same kinds of DMS studies. In particular, in Fig S4 we apply our same analysis pipeline for measured effects of replacements on protein stability (Δ*G* of folding as measured in (31)) rather than fitness or growth. We see that unlike the more integrated measures of protein function we have been examining so far, many of the corresponding distributions for protein stability effects have modes at intermediate values, consistent with the expectations based on physicochemical distances between amino acids. Overall, roughly half of the distributions are monotonic and are well approximated by the exponential law, though the shapes are more extreme than for DFEs, with few distributions close to uniform, and roughly 1/5 of the distributions exhibit intermediate modes, some of them quite distinctive (e.g., Thr to Asn, Ile to Val, Tyr to Phe). These results suggest that the truncated exponential law reflects something very general about mutational effects, applicable to measures of fitness across many different DMS studies, proteins, and protein functional assays, yet not applicable to these more basic biophysical measurements. We return to this issue in the Discussion.

Finally, it is worth noting that although we focus on modeling rank distributions here, transforming between a DFE for quantiles and a DFE on some other scale is straightforward. In particular, our maximum entropy distribution simply becomes the distribution that minimizes the Kullback-Leibler divergence from the overall DFE on the target scale to the type-specific DFE, subject to the constraint that a draw from the type-specific DFE is greater than the overall target DFE a specified fraction of the time. Practically, this translation to a new target measurement scale requires a parametric inverse cumulative distribution function marginalized over all types of mutations in the target scale (see Materials and Methods), enabling flexible modeling of pair-specific fitness effects in a wide variety of settings.

### A model of purifying selection recovers patterns of molecular evolution

Distributions of fitness effects from DMS experiments suggests that knowing the mean fitness effect for some type of amino acid change often provides little information: many distributions have a mean close to 0.5, and the dynamic range is only 3-fold, from a maximum of 0.71 to a minimum of 0.22. By contrast, molecular evolution exhibits much more distinctive patterns, as illustrated by the more than 10-fold dynamic range of Tang’s U (32), which measures how much more or less likely a non-synonymous mutation of a specific type is to go to fixation relative to a random mutation (33) based on computational analysis of natural sequence divergence.

Here, we attempt to reconcile these observations based on the assumption that in natural evolution the mutations that fix will be much fitter than random mutations, and therefore will be drawn primarily from the rightmost portion of the DFEs. Critically, when the DFEs have an exponential shape like the ones reported here, the exponential shape will tend to magnify the differences between exchange-specific DFEs with regard to both the highest and lowest fitness quantiles, e.g., a modest 1.5-fold difference in means corresponds to a more than a 3-fold difference in the chance of being in the top 10 % (Fig. 3a).

**Fig. 3.**
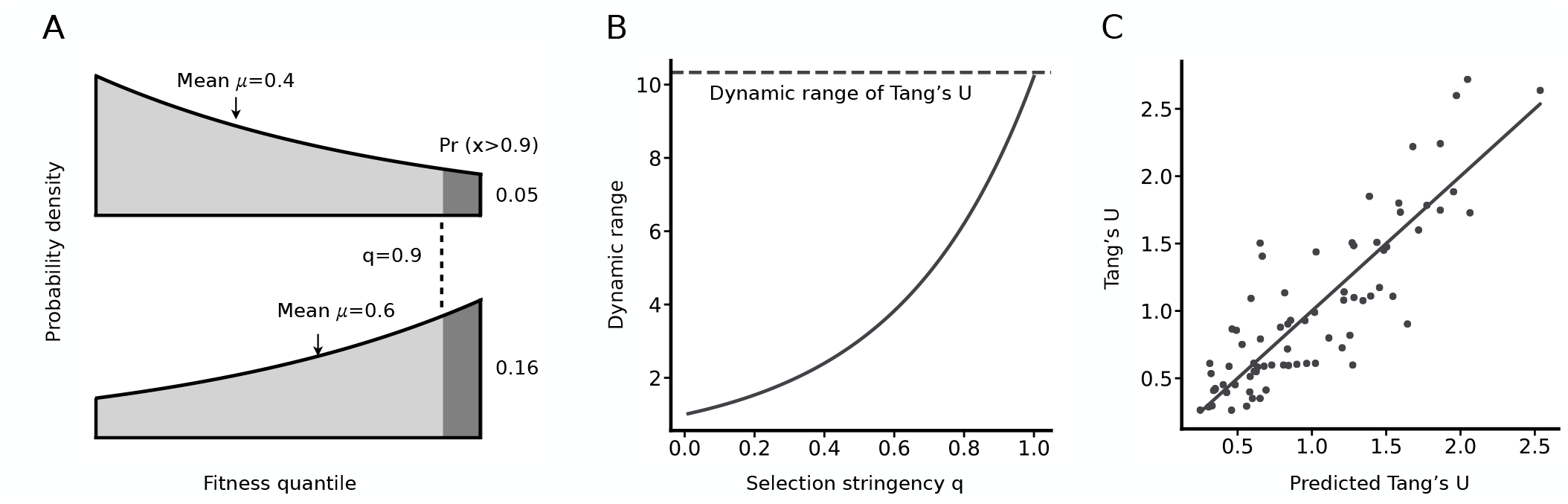
Selective filtering provides a quantitative model for patterns of evolution. For the subset of “singlet” replacements accessible via a single-nucleotide mutation, evolutionary exchangeability is represented by Tang’s *U*, which is derived from alignments of evolved sequences by a method designed to exclude mutational effects and to focus only on selection (32). (A) Given the truncated exponential model for DFEs, modest differences in the mean correspond to much larger differences in the top of the distribution, e.g., a difference of 0.4 vs. 0.6 in the mean corresponds to a 3-fold difference in the chance of exceeding a threshold *q* = 0.9. (B) Applying threshold selection to the DFEs for the singlet replacements shows that the dynamic range increases as *q →* 1, until it roughly matches the dynamic range of *U* (dashed line). (C) The predicted *U* matrix as *q →* 1 shows a good correlation with the observed Tang’s *U* (Pearson’s *r* = 0.86, *p* = 5 *×* 10^−23^), well described by the diagonal line *y* = *x*.

To more precisely formalize this hypothesis, suppose that we can approximate the effect of selection on protein evolution using a threshold *q* ranging from 0 to 1, such that amino acid replacements with a fitness quantile below *q* are rejected, and those above this threshold are accepted. Each value of *q* induces a different set of 380 frequencies of acceptable mutations, and the relative differences between these frequencies are expected to become more extreme as *q* approaches 1. In particular, under this model, if we write the normalized substitution rate for an amino acid change whose corresponding maximum entropy distribution has parameter *λ* as *K*_*λ*_*/K*_*s*_ (i.e. as a type-specific analog to *K*_*a*_*/K*_*s*_, or nonsynonymous/synonymous substitution rate ratio (34)), then as a function of *q* this normalized rate of evolution is given by

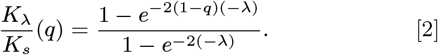

(The astute reader will recognize this expression as being formally identical to the Kimura probability of fixation (35) for a mutation with scaled selection coefficient −2*λ* and initial frequency 1 − *q* in a Wright-Fisher population, albeit obtained by an unrelated argument). As the selective stringency *q* increases towards 1, we then find that the ratio between *K*_*λ*_*/K*_*s*_ and *K*_*a*_*/K*_*s*_ = 1 − *q* converges to

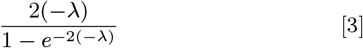

so that types of substitutions with negative *λ* are linearly enriched and types of substitutions with positive *λ* are exponentially depleted.

Prior studies of sequence divergence patterns provide some guidance for choosing reasonable values of *q*. In particular, values for *K*_*a*_*/K*_*s*_, the normalized rate ratio of amino acid changes to synonymous changes, are often in the range of 0.1 to 0.3 (36, 37). Under the simplifying assumption that all synonymous mutations are neutral, and that beneficial changes can be ignored, a *K*_*a*_*/K*_*s*_ value of 0.2 would mean that 80 % of non-synonymous mutational effects are deleterious, implying *q* = 0.8; moreover, if some fraction of synonymous mutations is deleterious, the implied value of *q* will be correspondingly higher.

To assess how well this kind of model recovers evolutionary patterns, we use the U matrix of “universal acceptability” from Tang, et al. (32) as our target of prediction. The U matrix, which applies to the 75 pairs of amino acids that are interchangeable by single-nucleotide mutations under the standard genetic code, aims to provide a pure measure of acceptability by selection: it is derived from amino acid sequence alignments by a method that controls for the genetic code, sequence composition, and mutation bias. Specifically, as shown by (33) the entries of the *U* matrix are proportional to *K*_*i*_*/K*_*s*_, the normalized rate of substitutions of the *i*-th of these 75 pairs of mutationally adjacent amino acids. In Materials and Methods, we show that these *K*_*i*_*/K*_*s*_ can be estimated as the harmonic mean of the *K*_*λ*_*/K*_*s*_ derived using the values of *λ* for the forward and backwards directions of the exchanges, so that given a value of selective stringency *q* we can use the values of *λ* inferred from deep mutational scanning experiments to estimate the entries of the U matrix. In particular, to the extent that our threshold model of selection on DFEs approximates acceptability in the natural evolutionary divergence of proteins, it is a predictor of U, and likewise, its correspondence with U is an independent measure of its accuracy as an evolutionary model of selective preferences.

Indeed, we find that, as *q* increases towards 1, the dynamic range of the predicted U values approaches the observed dynamic range of U (Fig. 3b) and the correlation between predicted and observed U values becomes strong (Pearson’s *r* = 0.86, *p* = 5 *×* 10^−23^), as shown in Fig. 3c. Note again that this predictor is derived entirely independently of Tang’s *U* : values of *U* are computed from large sets of alignments of naturally evolved protein sequences (32), whereas our predictor is computed by modeling selection as the left-truncation of empirical DFEs from DMS experiments, i.e., of the red curves in Fig. 2.

## Discussion

We used fitness quantiles from a large and diverse set of DMS studies to characterize DFEs (distributions of fitness effects) for the 380 types of amino acid changes in proteins. Many DFEs are nearly uniform, such that the identities of the starting and ending amino acids, by themselves, are not informative, reflecting the importance of molecular context in determining the effects of mutations. In general, the shapes of these DFEs are surprisingly well approximated by a truncated exponential distribution, corresponding to the maximum entropy distribution given the probability that a mutation of a given type is fitter than a random mutation. The shape of this maximum entropy distribution, in turn, suggests a resolution to the apparent contradiction between the predictable patterns of amino acid change observed in evolutionary sequence comparisons versus the difficulty of predicting mutational effects of SNPs or during protein engineering because the differences between exchanges are largest for the highest and lowest fitness quantiles. Specifically, we we show that a model of selective filtering where only the fittest mutations are allowed to fix applied to these maximum entropy DFEs closely reproduces the patterns of acceptability seen in evolutionary divergence.

Our maximum entropy model of DFEs is best understood as a model of conservatism rather than innovation, covering deleterious effects ranging from lethality to neutrality. The reason for this is that the fitness assays in DMS studies are designed by experimenters with a focus on wild-type functionality, so that beneficial variants are typically rare and small in effect. For the same reasons, the resulting model of evolutionary change is best understood as a model of conservatism or neutrality, not necessarily useful for modeling adaptive changes, which might be either conservative or radical– an open question in evolutionary biology (38, 39).

More generally, because the shape of the entire distribution follows if one knows the mean, and because the mean quantile can be estimated precisely from a modest amount of ordinal data (i.e., rankings), it should be possible to estimate distributions of exchangeability from much smaller and cruder sets of data than the one used here. This is relevant to the prospects for future experiments to explore effects of diversity and context. For instance, it has been suggested that evolutionary acceptability of transitions vs. transversions varies taxonomically (40): if the exponential model holds, exploring this type of hypothesis would require only a small amount of data on the fitness rankings of transitions and transversions in diverse taxa, because (when the exponential model holds) the shape of the high end follows precisely from the mean, which is easily estimated from a small amount of data.

Importantly, we find that the truncated exponential model is not universally applicable to mutational effects. It applies well to data from measurements of cellular fitness and protein function shown in Fig. 2, but provides a much poorer fit for the thermostability data shown in Fig. S3, where the distributions of quantiles often look more like the expectations from the chemical-similarity model (Fig. 1). In particular, the distributions are generally sharper, and many have intermediate modes, e.g., almost all changes from Gly to other amino acids in Fig. S3 have an intermediate mode. Presumably we see this difference because the distribution of thermostability effects is much less sensitive to context, i.e., a particular type of amino acid change can be understood as (for instance) the loss of an opportunity for a hydrogen bond or the addition of a methylene group, and this has a more consistent (less context-dependent) effect on thermostability than on protein function more generally, where folding stability will typically be only one of several different relevant properties.

The results reported here have many implications and potential applications in regard to modeling sequence evolution, the interpretation and modeling of variant effects in biomedical contexts, and in engineering and synthetic biology applications. Because mean quantile values only range from roughly 0.2 to 0.7, actual measurement data on the effects of amino acid changes do not support the common practice of offering token explanations of the form “this Arg to Lys change at position 127 is tolerated because Arg-to-Lys changes are conservative”. The problem with this type of claim is that an Arg-to-Lys change is only better than a random change 62 % of the time (upper left corner of Fig. 2). A more justifiable interpretation would be that, although Arg-to-Lys changes are not intrinsically benign, they are several times more likely than random changes to be in the top 10 % most benign mutations.

By contrast to the relative lack of informativeness of mean values applied to specific outcomes, the tunable model of selective stringency is potentially highly informative if one is optimizing or making inferences over a large set of outcomes. For instance, in a synthetic biology context, if one is constructing a finite library to explore the sequence space within *n* changes of the starting sequence (where *n* is a small number like 3 or 8), weighting the options by the chance of being in the top decile of fitness may provide a considerable improvement over a naive randomization, and the DFEs provided here are a better choice to guide optimization than amino acid scoring matrices (e.g., BLOSUM), which are not pure measures of functional effect, but reflect both mutational biases and the genetic code (see 41, 42).

An even more promising approach would be to integrate a model of selective filtering into a generalized maximum likelihood approach to phylogenetic inference such as IQ-TREE (43), such that different values of *q* could be inferred for different branches of the tree or for different proteins. A relatively clear prediction from basic population genetics is that the inferred *q* will be higher in larger populations where selection is stronger relative to drift. Weber and Whelan (44) report that, in larger populations, biochemical properties are more strongly predictive for patterns of amino acid change, suggesting that this kind of effect is strong enough to be observed in available data. Thus it may be possible to infer historical changes in the strength of selection (and by inference, e.g., population size) solely from patterns of relative amino acid exchangeability.

## Materials and Methods

### Data Collection and Preprocessing

We acquired the ProteinGym v0.1 dataset from https://github.com/OATML-Markslab/Tranception. This dataset includes measurements of effects for various mutations, including some that change multiple amino acids. We pruned this data set to include solely on single amino-acid changes, resulting in mutational effects for 358,080 replacements from 87 assays. For within-assay mutation ranking, we employed the percentile ranking function from Python’s pandas package, using the ‘average’ method to break ties. Additionally, to compare mutation effects from experimental data with substitution patterns in molecular evolution, we sourced the values of Tang’s U from Table 2 in (32). Finally, we downloaded a data set of mutational effects on protein stability(45) from https://zenodo.org/records/7992926. This dataset includes measurements of the effects of 389,068 single amino-acid mutations across 412 proteins on protein folding stability. The preprocessing method follows the same approach as our handling of the ProteinGym data, where raw measurements are converted into percentile rankings within each protein.

### Fitting Truncated Exponential Distributions from Sample Mean

The probability density function for a truncated exponential distribution, as described, takes the form *p*(*x*) = *ce*^−*λx*^,where the distribution is bounded between 0 and 1 and 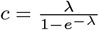 serves as the normalizing constant. Here, *λ* is a positive real number. The expected value, or mean, of this distribution is given by:

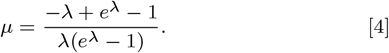

This function is known to be monotonic (27). Consequently, the sample mean, *μ*, uniquely determines *λ*. However, there is no closed-form expression to derive *λ* from *μ*, necessitating a numerical approach. We employ Newton’s method to calculate *λ* from the sample mean, *μ*. For our analysis, we have utilized the complete set of rank-transformed mutation effect data to estimate the sample mean *μ* for each type of amino-acid mutation unless otherwise indicated.

### Theoretical Mean-Variance Relationship in Truncated Exponential Distributions

The variance of a truncated exponential distribution with parameter *λ* is given analytically by:

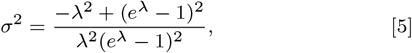

and thus the relationship between the variance *σ*^2^ and the mean *μ* can be depicted as a parametric curve in terms of *λ*. As shown in Fig. 3a, this curve displays an inverted U shape and is symmetric about the line *x* = 0.5.The curve peaks when *μ* = 0.5, at which point the truncated exponential distribution becomes a uniform distribution.

### Converting Rank DFE to DFE in other scales

Given a type-specific rank DFE specified by equation 1, denote the inverse cumulative distribution function (CDF) of the marginal distributions over all types of mutations in another scale as *F* ^−1^(*x*). The inverse CDF *G*^−1^(*x*) for the focal type-specific DFE in the target scale is the composite function of *F* ^−1^(*x*) and the inverse CDF of the truncated exponential rank distribution function:

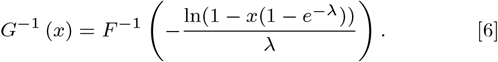

Noting that by definition the distribution of the overall DFE on our percentile-rank scale is the uniform distribution on [0,1], we see that the Kullback-Leibler divergence from this distribution to any other distribution on the percentile-rank scale is the negative of the entropy. Because the Kullback-Leibler divergence is invariant under changes of scale (invertible transformations (27)), and *F* and *G* differ only by a change of scale from the overall DFE and the maximum-entropy type-specific DFE on our percentile-rank scale, the fact that our type-specific distribution maximizes the entropy given the probability that a mutation of that type is fitter than a random mutation shows that the distribution with CDF *G* above minimizes the Kullback-Leibler divergence from the distribution with CDF *F* under this same constraint.

### Linking Tang’s U to parameters of Experimental DFEs

Conceptually, Tang’s *U* index (32) for a particular pair of amino acids aims to measure the fixation probability between that particular pair of amino-acids, relative to the fixation probability for a random amino acid exchange (33). Following (33), we can consider a codon model in which the rate of evolution from codon *i* to codon *j* ≠ *i* is given by:

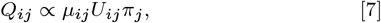

where *μ*_*ij*_ is a symmetric mutational bias (*μ*_*ij*_ = *μ*_*ji*_), *U*_*ij*_ is the entry of the *U* matrix corresponding to the amino acids encoded by *i* and *j* or is 1 if the mutation is synonymous (*U* is also symmetric, *U*_*ij*_ = *U*_*ji*_), *π*_*j*_ is the stationary frequency of codon *j*, and the diagonal entries of *Q* are chosen so that the row sums all equal 0. It is easy to show that the continuous time Markov chain with rate matrix *Q* has the stationary distribution specified by *π*, and moreover that *Q*_*ij*_*/Q*_*ji*_ = *π*_*j*_*/pi*_*i*_ so that solving for *U*_*ij*_ we have *U*_*ij*_ ∝2*/* ((1*/Q*_*ij*_) + (1*/Q*_*ji*_)). Substituting in *Q*_*ij*_ ∝ *K*_*λ*_*/K*_*s*_(*q*) under the selective stringency model into this harmonic mean formula gives the estimate of the corresponding entry in Tang’s U matrix described in the main text, where we have centered the predicted values of *U* so that the mean across the 75 exchanges is 1 to match the published scaling convention for Tang’s U (32).

### Availability of data and scripts

The Python code and raw data files for generating all figures and performing statistical analyses as described in the manuscript have been deposited in a GitHub repository: https://github.com/mengysun/AA_exchangeability.

## ACKNOWLEDGMENTS

Funding for this work was provided in part by NIH grant R35 GM133613 (D.M.M.), the John Templeton Foundation (grant #61782, D.M.M.), and additional funding from the Simons Center for Quantitative Biology at Cold Spring Harbor Laboratory. The identification of any specific commercial products is for the purpose of specifying a protocol, and does not imply a recommendation or endorsement by the National Institute of Standards and Technology.

## Author Contributions

All authors contributed to the experimental design; MS carried out the data analysis and calculations, and drafted a manuscript; MS, AS and DMM revised the manuscript.

## Author Declaration

The authors declare that they have no conflicts of interest.

## Supporting information

**Fig. S1.**
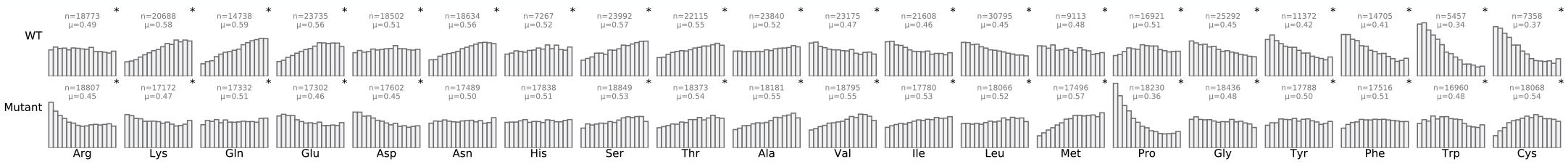
Quantile distributions of fitness effects from ProteinGym, categorized by wild-type (“from”) and mutant (“to”) amino acids. The distributions in the top row aggregate all 358,080 mutational effects by the wild-tpe amino acid, and those in the bottom row aggregate the same mutational effects by the mutant amino acid. Distributions that are significantly different from a uniform distribution are marked with a star in the upper-right corner. The sample size for each distribution is indicated by *n*, and the mean of each distribution is indicated by *μ*.

**Fig. S2.**
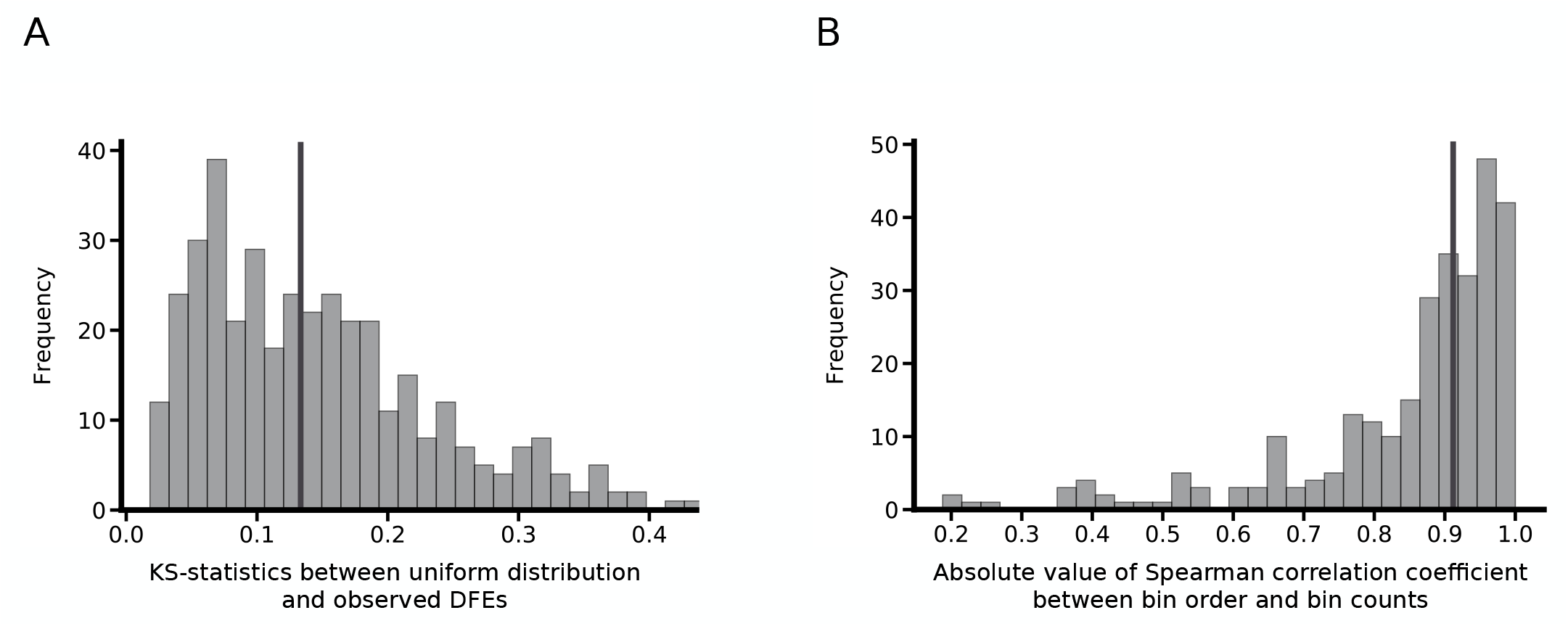
Statistical characteristics of 380 distributions of fitness effects. (A) Most of the distributions are broad, characterized by relatively small deviations from uniform distributions measured by the KS-statistics (vertical line indicates the median). (B) Most of the distributions tend to be monotonic, indicated by the highly right-skewed Spearman correlation coefficients between the bin order and bin counts (vertical line indicates the median).

**Fig. S3.**
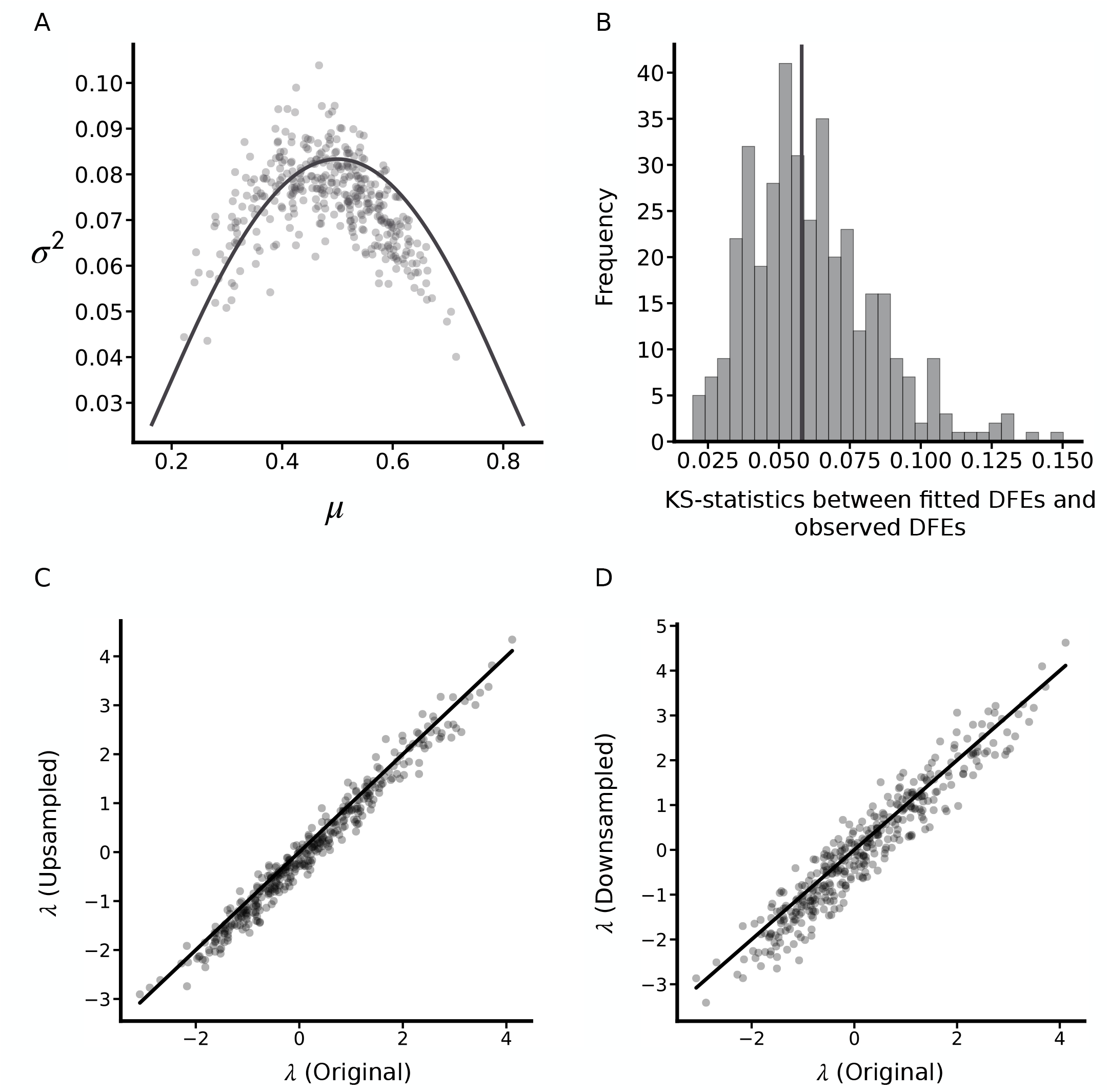
Truncated exponential captures the 380 DFEs well. (A) The theoretical relationship (solid line) between expectation and variance of truncated exponential distributions qualitatively matches the observed data. (B) The KS-statistics between fitted truncated exponential distributions versus individual DFEs tend to be very small (vertical red line indicates the median), suggesting that truncated exponential distribution provides a good fit to the data. (C) and (D): the *λ* values estimated by equalizing the representation of each type, either by upsampling to the maximum count of all mutation types in the original dataset (C), or by downsampling to the minimum (D), is highly correlated to the original estimation, indicating that the truncated exponential distribution observed is not an artifact of unequal distribution of amino-acids in natural proteins (solid line indicates *y* = *x*).

**Fig. S4.**
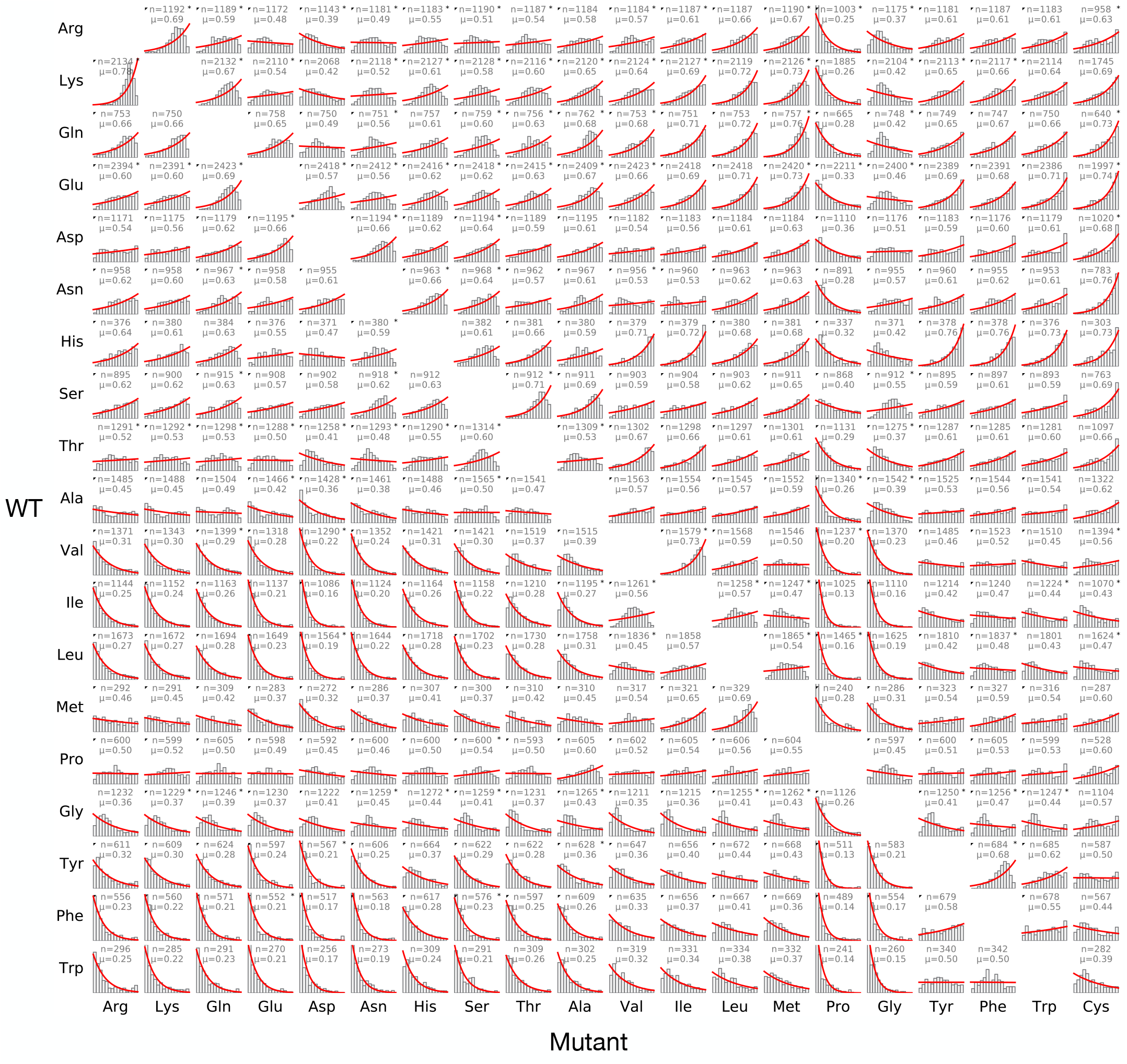
Distributions of mutational effects on protein-folding for 389,030 amino-acid-altering mutations, categorized by the wild-type (row) and mutant (column) amino acids. Histograms illustrate the observed fitness effect distributions for mutations relative to each amino acid type, along with the truncated exponential fit for each distribution (red lines). Distributions that significantly deviate from a truncated exponential distribution are marked with a star at the right corner, whereas black triangles in the upper left corner indicate the pairs with significant forward-reverse asymmetry (Bonferroni-corrected *p <* 0.05, two-sample Kolmogorov–Smirnov test). For each distribution, the mean, which is equal to the probability that a replacement of this type is fitter than a random mutation, is given by *μ*, while the number of observations is given by *n*.

## Bibliography

1. E Zuckerkandl, L Pauling, Evolutionary Divergence and Convergence in Proteins, eds. V Bryson, H Vogel (Academic Press, New York), (1965).

2. R Eck, M Dayhoff, Atlas of Protein Sequence and Structure. (National Biomedical Research Foundation, Silver Spring, MD), (1966).

3. S Kawashima, M Kanehisa, AAindex: amino acid index database. Nucleic Acids Res 28, 374. (2000).

4. P Sneath, Relations between chemical structures and biological activity in peptides. J. Theor. Biol. 12, 157 (1966).

5. CJ Epstein, Non-randomness of amino-acid changes in the evolution of homologous proteins. Nature 215, 355–9 (1967).

6. R Grantham, Amino acid difference formula to help explain protein evolution. Science 185, 862–864 (1974).

7. T Miyata, S Miyazawa, T Yasunaga, Two types of amino acid substitutions in protein evolution. J Mol Evol 12, 219–36 (1979).

8. SF Altschul, W Gish, W Miller, EW Myers, DJ Lipman, Basic local alignment search tool. J Mol Biol 215, 403–10 (1990).

9. R Durbin, SR Eddy, A Krogh, G Mitchison, Biological Sequence Analysis: Probabilistic Models of Proteins and Nucleic Acids. (Cambridge University Press, Cambridge), (1998).

10. JD Thompson, DG Higgins, TJ Gibson, CLUSTAL W: improving the sensitivity of progressive multiple sequence alignment through sequence weighting, position-specific gap penalties and weight matrix choice. Nucleic Acids Res 22, 4673–80 (1994).

11. LA Miosge, et al., Comparison of predicted and actual consequences of missense mutations. Proc Natl Acad Sci U S A 112, E5189–98 (2015).

12. BJ Livesey, JA Marsh, Using deep mutational scanning to benchmark variant effect predictors and identify disease mutations. Mol. systems biology 16, e9380 (2020).

13. BJ Livesey, JA Marsh, Updated benchmarking of variant effect predictors using deep mutational scanning. Mol. systems biology 19, e11474 (2023).

14. Z Wang, J Moult, SNPs, protein structure, and disease. Hum Mutat 17, 263–70 (2001).

15. J Domingo, P Baeza-Centurion, B Lehner, The Causes and Consequences of Genetic Interactions (Epistasis). Annu. Rev Genomics Hum Genet. 20, 433–460 (2019).

16. JF Storz, Compensatory mutations and epistasis for protein function. Curr Opin Struct Biol 50, 18–25 (2017).

17. Z Wu, SBJ Kan, RD Lewis, BJ Wittmann, FH Arnold, Machine learning-assisted directed protein evolution with combinatorial libraries. Proc Natl Acad Sci U S A 116, 8852–8858 (2019).

18. TN Starr, JM Flynn, P Mishra, DNA Bolon, JW Thornton, Pervasive contingency and entrenchment in a billion years of Hsp90 evolution. Proc Natl Acad Sci U S A 115, 4453–4458 (2018).

19. VO Pokusaeva, et al., An experimental assay of the interactions of amino acids from orthologous sequences shaping a complex fitness landscape. PLoS Genet. 15, e1008079 (2019).

20. VE Gray, RJ Hause, DM Fowler, Analysis of Large-Scale Mutagenesis Data To Assess the Impact of Single Amino Acid Substitutions. Genetics 207, 53–61 (2017).

21. VE Gray, RJ Hause, J Luebeck, J Shendure, DM Fowler, Quantitative Missense Variant Effect Prediction Using Large-Scale Mutagenesis Data. Cell Syst 6, 116–124 e3 (2018).

22. DM Fowler, S Fields, Deep mutational scanning: a new style of protein science. Nat Methods 11, 801–7 (2014).

23. JB Kinney, DM McCandlish, Massively Parallel Assays and Quantitative Sequence-Function Relationships. Annu. Rev Genomics Hum Genet. 20, 99–127 (2019).

24. P Notin, et al., Tranception: protein fitness prediction with autoregressive transformers and inference-time retrieval in 39th International Conference on Machine Learning. Vol. Proceedings of the 39th International Confjaynaerence on Machine Learning, (year?).

25. ET Jaynes, Information Theory and Statistical Mechanics. Phys. Rev. 106, 620–630 (1957).

26. ET Jaynes, Information Theory and Statistical Mechanics. II. Phys. Rev. 108, 171–190 (1957).

27. TM Cover, JA Thomas, Elements of Information Theory. (John Wiley and Sons, Inc., Hoboken, NJ), (2005).

28. O Cotto, T Day, A null model for the distribution of fitness effects of mutations. Proc Natl Acad Sci U S A 120, e2218200120 (2023).

29. MM Johnson, CO Wilke, Site-Specific Amino Acid Distributions Follow a Universal Shape. J Mol Evol 88, 731–741 (2020).

30. TC Day, et al., Cellular organization in lab-evolved and extant multicellular species obeys a maximum entropy law. Elife 11, e72707 (2022).

31. K Tsuboyama, et al., Mega-scale experimental analysis of protein folding stability in biology and design. Nature 620, 434–444 (2023).

32. H Tang, GJ Wyckoff, J Lu, CI Wu, A universal evolutionary index for amino acid changes. Mol Biol Evol 21, 1548–56 (2004).

33. H Tang, CI Wu, A new method for estimating nonsynonymous substitutions and its applications to detecting positive selection. Mol Biol Evol 23, 372–9 (2006).

34. M Nei, T Gojobori, Simple methods for estimating the numbers of synonymous and nonsynonymous nucleotide substitutions. Mol. Biol. Evol. pp. 418–426 (1986).

35. M Kimura, On the probability of fixation of mutant genes in a population. Genetics 47, 713–9 (1962).

36. D Wang, et al., How do variable substitution rates influence Ka and Ks calculations? Genomics Proteomics Bioinforma. 7, 116–27 (2009).

37. JB Wolf, A Kunstner, K Nam, M Jakobsson, H Ellegren, Nonlinear dynamics of nonsynonymous (dN) and synonymous (dS) substitution rates affects inference of selection. Genome Biol Evol 1, 308–19 (2009).

38. Q Chen, et al., Molecular evolution in large steps - Codon substitutions under positive selection. Mol Biol Evol (2019).

39. Q Chen, A Lan, X Shen, CI Wu, Molecular evolution in small steps under prevailing negative selection - A nearly-universal rule of codon substitution. Genome Biol Evol 11, 2702–2712 (2019).

40. Z Zou, J Zhang, Are nonsynonymous transversions generally more deleterious than nonsynonymous transitions? Mol Biol Evol (2020).

41. PC Ng, S Henikoff, Predicting deleterious amino acid substitutions. Genome Res 11, 863–74 (2001).

42. LY Yampolsky, A Stoltzfus, The exchangeability of amino acids in proteins. Genetics 170, 1459–1472 (2005).

43. LT Nguyen, HA Schmidt, A von Haeseler, BQ Minh, IQ-TREE: a fast and effective stochastic algorithm for estimating maximum-likelihood phylogenies. Mol Biol Evol 32, 268–74 (2015).

44. CC Weber, S Whelan, Physicochemical Amino Acid Properties Better Describe Substitution Rates in Large Populations. Mol Biol Evol 36, 679–690 (2019).

45. K Tsuboyama, et al., Mega-scale experimental analysis of protein folding stability in biology and design. Nature 620, 434–444 (2023).

